# Global Impacts of Climate Change on Avian Functional Diversity

**DOI:** 10.1101/2020.06.01.127779

**Authors:** Peter S. Stewart, Alke Voskamp, Matthias F. Biber, Christian Hof, Stephen G. Willis, Joseph A. Tobias

## Abstract

Climate change is predicted to drive geographical range shifts in many taxa, leading to the formation of novel species assemblages and fluctuations in species richness worldwide. However, the effect of these changes on functional diversity is not yet fully understood, in part because comprehensive species-level trait data are generally lacking at global scales. Here we use morphometric and ecological trait data for 8269 terrestrial bird species to compare functional diversity (FD) of current and future bird assemblages under a medium emissions scenario. We show that future assemblages are likely to undergo substantial shifts in trait structure, with the direction and magnitude of these shifts varying with geographical location and trophic guild. Specifically, invertivore FD is projected to increase at higher latitudes with concurrent losses at mid-latitudes, reflecting poleward shifts in range, whereas frugivore FD is projected to fluctuate in many tropical regions with major declines in much of South America and New Guinea. We show that these projected changes in FD are generally greater than expected from changing species richness alone, indicating that projected FD changes are primarily driven by the loss or gain of functionally distinct species. Our findings suggest that climate change will drive continental-scale shifts in avian functional diversity, with potentially far-reaching implications for ecosystem functions and resilience.

## Introduction

Climate change is driving substantial shifts in species distributions worldwide (Walther et al., 2002; Parmesan & Yohe, 2003). The magnitude and direction of these shifts vary across species, generating novel species assemblages which differ in structure and composition to those observed today (Williams & Jackson, 2007), potentially resulting in changes to ecosystem functions, services and resilience (Barbet-Massin & Jetz, 2015). Understanding and forecasting these changes to assemblage structure is an important step towards developing effective conservation strategies targeted at regions where ecosystem functions are likely to be compromised by climate change (Oliver & Roy, 2015). However, few previous studies have gone beyond relatively simplistic estimates of changes in species distributions (Barbet-Massin & Jetz, 2015; Gaüzère, Jiguet, & Devictor, 2015), and thus the likely trophic structure and functioning of future assemblages under climate change remains unclear.

One method for estimating whether the functioning of future ecosystems is likely to be altered or impaired is to assess changes in the functional traits present in species assemblages (Barbet-Massin & Jetz, 2015). Functional traits provide information about key dimensions of the ecological niche (Winemiller, Fitzgerald, Bower, & Pianka, 2015; Pigot et al., 2016; Pigot et al., 2020), and functional diversity (FD) quantifies the range of functional traits present within a community (Petchey & Gaston, 2002). Projected shifts in FD can therefore provide information about changes in the diversity and characteristics of niches occupied within species assemblages (Mouillot, Graham, Villéger, Mason, & Bellwood, 2013; Pigot, Trisos, & Tobias, 2016), and thus the likelihood of assemblages to sustain important ecological processes (Leitão et al., 2016; Petchey & Gaston, 2002; Tilman, 2001; Villéger, Mason, & Mouillot, 2008). Specifically, a reduction in trait diversity or a shift in trait structure suggests ecosystem functions will be lost or altered (Tilman et al., 1997; Cardinale, Palmer & Collins, 2002; Díaz et al., 2013), potentially impacting ecosystem multifunctionality (Mouillot et al., 2011), and resilience (Bregman et al., 2016).

Assessing these functional impacts at macroecological scales has proved challenging because detailed trait data are generally available only patchily across large numbers of species. Most previous research has therefore focused at smaller spatial or taxonomic scales (e.g. Biswas, Vogt, & Sharma, 2017; Van Zuiden et al., 2016; Mokany et al., 2015; Gaüzère, Jiguet, & Devictor, 2015), making it difficult to know how far their results can be generalised. The only datasets available at a global scale that have previously been used to assess climate change impacts on assemblage trait structure are largely based on relatively crude species traits, such as diet categories or binary characters (e.g. diurnal versus nocturnal) (Barbet-Massin & Jetz, 2015). The main drawback of these categorical traits is that vital information is lost during the process of categorisation, with many distinctly different species lumped within each particular category (Laureto, Cianciaruso, & Samia, 2015; Weiher et al., 1999).

To address this issue, we compared the functional structure of current and future avian assemblages using a comprehensive dataset of morphological and ecological traits for 8269 terrestrial bird species worldwide (Pigot et al., 2020). For all species, we compiled eight continuous variables to capture variation in beak shape, wing shape, and the length of tarsus and tail. Together, these morphological traits provide and index of avian dispersal ability and trophic niche (Claramunt et al., 2012; Tobias et al., 2014; Pigot et al., 2020). In contrast, previous global studies have mainly been limited to a single continuous ecological trait – body mass – which is a poor predictor of avian dispersal (Sheard et al. 2020) and only weakly informative about ecological niche differences (Pigot et al., 2020). We used this comprehensive trait dataset to calculate the functional diversity of future species assemblages estimated using recent species range projections (Hof et al., 2018) with a time horizon centred on 2050. Finally, we included additional analyses partitioning avian assemblages into two major dietary groups – frugivores and invertivores – which play important ecological roles through mutualistic interactions and insect predation, respectively (Pigot et al., 2016; Pigot et al., 2020; Şekercioǧlu, 2006).

Specialist avian frugivores are vital seed dispersal agents, especially for large-fruited or large-seeded plants in tropical regions (Corlett, 2017; Snow, 1981). The loss of avian frugivores can substantially reduce recruitment of juvenile plants and limits the ability of plants to track climate change (Cordeiro & Howe, 2003; McConkey et al., 2011; Mokany, Prasad, & Westcott, 2014). Ultimately, declines in frugivore diversity may therefore impair carbon storage, potentially driving further climatic change (Bello et al., 2015). Avian invertivores in terrestrial ecosystems exert top-down control on invertebrate populations, including numerous phytophagous (herbivorous) insects and their larvae, thereby indirectly benefitting plant populations (Mäntylä, Klemola, & Laaksonen, 2011) and boosting ecosystem productivity (Marquis & Whelan, 1994). Invertivores also provide an important ecosystem service by limiting the impact of insect pests on agricultural crops (Jones, Sieving, & Jacobson, 2005; Karp et al., 2013; Mols & Visser, 2002).

The main aims of our study are to (1) re-evaluate the future impacts of climate change on the FD of avian assemblages, with particular focus on the structure of frugivore and invertivore communities, and (2) quantify the relative turnover of functionally redundant and functionally distinct species to determine whether changes in FD are primarily driven by the loss or gain of functionally distinct species. This second aim is related to the findings of Barbet-Massin and Jetz (2015), who highlighted the importance of functional turnover – the replacement of one subset of species by another set of species – in determining the impact of climate-driven range shifts on FD. They reported that projected shifts in FD were smaller than expected from observed changes in avian species richness, suggesting a tendency for turnover of functionally redundant species which contribute relatively little to the overall FD of the assemblage. We revisit this question in the context of continuous traits to examine the role of turnover in explaining projected shifts in FD for frugivorous and invertivorous birds.

## Materials and Methods

### 1. Current and Projected Avian Distributions

We used baseline and projected distributions for 8269 terrestrial bird species worldwide from Hof et al. (2018) as binary presence-absence matrices, where each row represented a 0.5° × 0.5° latitude-longitude grid cell (henceforth, “grid-cell assemblage”), and each column a bird species (1 = present, 0 = absent). The presence-absence data were derived by thresholding the projected probabilities of occurrence, using true skill statistics (TSS; Allouche, Tsoar & Kadmon, 2006). As detailed in Hof et al., (2018), baseline distributions were derived from expert range maps produced by BirdLife International and NatureServe (2015), considering only areas where a species was extant or probably extant, was a native species and was resident or occurred regularly during the breeding season. Antarctica was excluded, leaving a total of 65546 grid-cell assemblages.

Species distribution models were used by Hof et al. (2018) to generate projected distributions for each bird species. These used current (1980-2009) data on temperature and precipitation from the meteorological forcing dataset EWEMBI (Lange, 2016) and future data from bias-corrected global climate scenarios produced by ISIMIP phase 2b (Frieler et al., 2017) for the year 2050 under the representative concentration pathway RCP6.0 (Fujino et al., 2006; Hijioka et al., 2008), incorporating four different global climate models (GCMs, GFDL-ESM2M, HadGEM2-ES, IPSL-CM5A-LR and MIROC5) which were chosen to reflect a representative spread of responses among the many GCMs available. We chose to use future projections for the period centred on 2050 because this period is most relevant for current climate change policy timelines and for conservation and ecosystem service mitigation planning. In addition, uncertainty in projections of biological change typically increase markedly for end-of-century projections (Baker et al. 2015). We selected RCP6.0 as a realistic medium-high climate-policy intervention scenario, which assumes a shift from coal towards oil, gas and renewables for energy (Masui et al. 2011). Two types of species distribution model (a general additive model, and a generalized boosted regression model) under each of the four global climate models were used to derive an ensemble projection. The models used pseudo-absences selected using a distance-weighted approach, in which absences beyond each species’ baseline distribution were selected following a declining probability of 1/(D_e_)^2^ where D_e_ is distance from the edge of the distribution (Hof et al., 2018). An intermediate dispersal scenario was used in which species could disperse to grid cells within d/2 of their baseline range (d = baseline range diameter) and were constrained to neighbouring zoogeographic realms (Hof et al., 2018). We chose to use only a single dispersal scenario because while choice of dispersal range does affect the magnitude of species richness changes, particularly for species-poor assemblages, it has little influence on the broad-scale patterns observed (Hof et al., 2018). A blocking approach, as per Bagchi et al. (2013), was used to reduce the effect of spatial autocorrelation. For species with too few occurrences to perform blocking, models were instead on a random subset of 70% of the data and performance was evaluated using the left-out 30%. Model validation was performed using the ‘area under curve’ (AUC) method to assess how models performed when used to predict current bird ranges (Hof et al., 2018). Species for which models performed poorly (AUC<0.7 across all models) were not included in the baseline or for future projections. Furthermore, only species which were terrestrial and which occupied 10 or more grid cells were included due to the difficulties associated with modelling species for which there are few presences (Hof et al., 2018). Therefore, of the 9882 bird species for which baseline data are available, 8269 species were considered in our analyses.

### 2. Functional Traits

For our species sample (n = 8269), we assembled morphological and ecological trait data from a comprehensive global dataset (Pigot et al., 2020). For each species, we compiled information on eight continuous traits (see Table S1) which provide information about key dimensions of the avian niche. Specifically, beak is linked to the trophic niche by indicating the size and type of food consumed (Lederer, 1975; Wheelwright, 1985; Hsu, Shaner, Chang, Ke, & Kao, 2014; Pigot et al., 2020). Tarsus length, wing chord, first-secondary length and tail length are locomotory traits related to microhabitat utilisation, foraging strategy, and dispersal (Claramunt et al., 2012; Miles & Ricklefs, 1984; Miles, Ricklefs, & Travis, 1987; Sheard et al., 2020). Finally, body mass is related to various aspects of the avian niche including metabolic requirements (McGill et al., 2006), movement (Wotton & Kelly, 2012), and foraging behaviour (Dial, Greene, & Irschick, 2008). Beak length was taken from the tip to the anterior nares, as this metric has a lower measurement error than measuring from the base of the skull (Borras, Pascual, & Senar, 2000). In combination, these traits provide an eight-dimensional quantitative morphological space⎯hereafter termed “morphospace”⎯in which the position of each species reflects key aspects of the trophic niche, including trophic level, diet and foraging behaviour (Pigot et al. 2020).

All trait values were log-transformed then z-transformed and converted into a distance matrix with the package Cluster (Maechler *et al.*, 2018). We further adapted the matrix into a nested functional dendrogram using the “average” clustering algorithm in the ape package (Paradis and Schliep, 2018). Additionally, we combined the z-transformed traits in a principal component analysis (PCA). We only retained the first four principal component (PC) axes for further analysis because the computational requirements of functional diversity indices increase rapidly with increasing dimensionality and a four-dimensional morphospace is sufficient to describe variation in avian trophic niches (Pigot et al., 2020). Together, these axes accounted for >95% of the variance in the functional trait data (see Table S2 for PC scores). All analyses were performed in R v3.5.1 (R Core Team, 2018).

Analysing functional trait structure of entire assemblages combines information across multiple different trophic levels and niches. While this provides a useful overview of general patterns, it can mask effects specific to particular ecological processes, unless these processes are partitioned into separate analyses (Bregman et al. 2014; Bregman et al. 2016; Cannon et al. 2019). To address this issue, we used published datasets (Tobias & Pigot, 2019; Pigot et al., 2020) to subdivide our species sample into those classified as frugivores and terrestrial invertivores, which are associated with important ecosystem functions and services, namely seed dispersal and the top-down control of invertebrates, respectively (Şekercioǧlu, 2006; Karp *et al.*, 2013; Bregman et al., 2016). Frugivores and terrestrial invertivores were defined as those terrestrial (i.e. non-aquatic) species for which >50% of the diet consists of fruit or invertebrates, respectively.

### 3. Analysis of Trait Structure

We used current and future geographical ranges for 8269 bird species to calculate species richness occurring in all grid cells worldwide, producing a dataset of 65546 pairs of current and future assemblages. To quantify the trait structure of each grid-cell assemblage, we calculated four functional diversity metrics: Functional Diversity (FD; Petchey & Gaston, 2002), Functional Richness (FRic; Villéger, Mason & Mouillot, 2008), and Gaussian hypervolume (Blonder, 2017) volume (Hvol) and centroid coordinates (Hcent).

FD, the sum of branch lengths on a functional dendrogram, measures how species are dispersed in trait space (Petchey & Gaston, 2002), with greater values of FD indicating a greater degree of trait complementarity. We calculated FD from the functional trait dendrogram for each assemblage, using the R package ape (Paradis and Schliep, 2018).

FRic, the volume of the smallest convex hull enclosing all trait values in an assemblage (Villéger et al., 2008), does not measure ‘functional richness’ per se because it is entirely dependent on the species with the most extreme trait values. However, it provides a useful indicator of the gain or loss of functionally extreme species. We calculated FRic based on the trait PC scores for each presence-absence-matrix row in all assemblages with five or more species using R package geometry (Habel *et al.*, 2015).

The volume of a Gaussian hypervolume (Hvol) provides a different perspective to dendrogram-based approaches in that it focuses on the volume of morphospace occupied by species in an assemblage (Maire, Grenouillet, Brosse, & Villéger, 2015; Blonder, 2017). A final method focuses on movement of the hypervolume centroid (Hcent), which can indicate important shifts in trait structure even in the absence of a change in functional diversity. Hvol and Hcent were calculated for each assemblage from the trait PC scores using the Silverman bandwidth estimate in the R package hypervolume (Blonder and Harris, 2018).

To measure how trait structure differed between current and future assemblages, the difference in FD, FRic and Hvol values between the baseline and projected distributions (ΔFD, ΔFRic, and ΔHvol) were calculated for each assemblage, as well as the Euclidean distance between the baseline and projected Hcent coordinates. Projected changes in species richness (ΔSR) were also calculated for each assemblage. Using the coordinates of each grid-cell assemblage, we plotted maps of each metric (ΔFD, ΔFRic, ΔHvol, Hcent Euclidean distance, and ΔSR) to examine the geographical distribution of our projected changes.

To examine whether changes in FD were primarily driven by the gain or loss of functionally distinct species, observed ΔFD was compared to expected ΔFD values predicted from ΔSR. To reduce sensitivity to outliers, we used robust regression (Fox & Weisberg, 2012), using the R package MASS (Venables and Ripley, 2002) with ΔFD as the response variable and ΔSR as the explanatory variable. The model output was used to predict ΔFD from ΔSR, and the difference between observed and predicted ΔFD was calculated (Residual FD). We then categorized each assemblage based on the percentage difference between observed and predicted ΔFD (=residual/predicted). Assemblages in which FD was lost despite species richness (SR) gain (and vice versa) and where FD changed despite no SR change were given separate categories.

We conducted cladewide analyses at a global level for all 8269 species, and then repeated analyses separately for frugivores (n = 898 species) and invertivores (n = 4347 species).

## Results

### a. Cladewide analyses

In our geographical range forecasts, 1.1% of bird species (n = 89) had no suitable projected climate within their dispersal buffer and were thus considered to become extinct by 2050. Shifts in the geographical distribution of the remaining 98.9% of bird species altered the diversity and structure of most grid-cell assemblages. The change in species richness between current (baseline) and future (projected) assemblages varied by geographic region (Figure 1a, Figure S1a), ranging from losses of 286 species to gains of 387 species (mean = 3.281, sd = 31.3).

**Figure 1.**
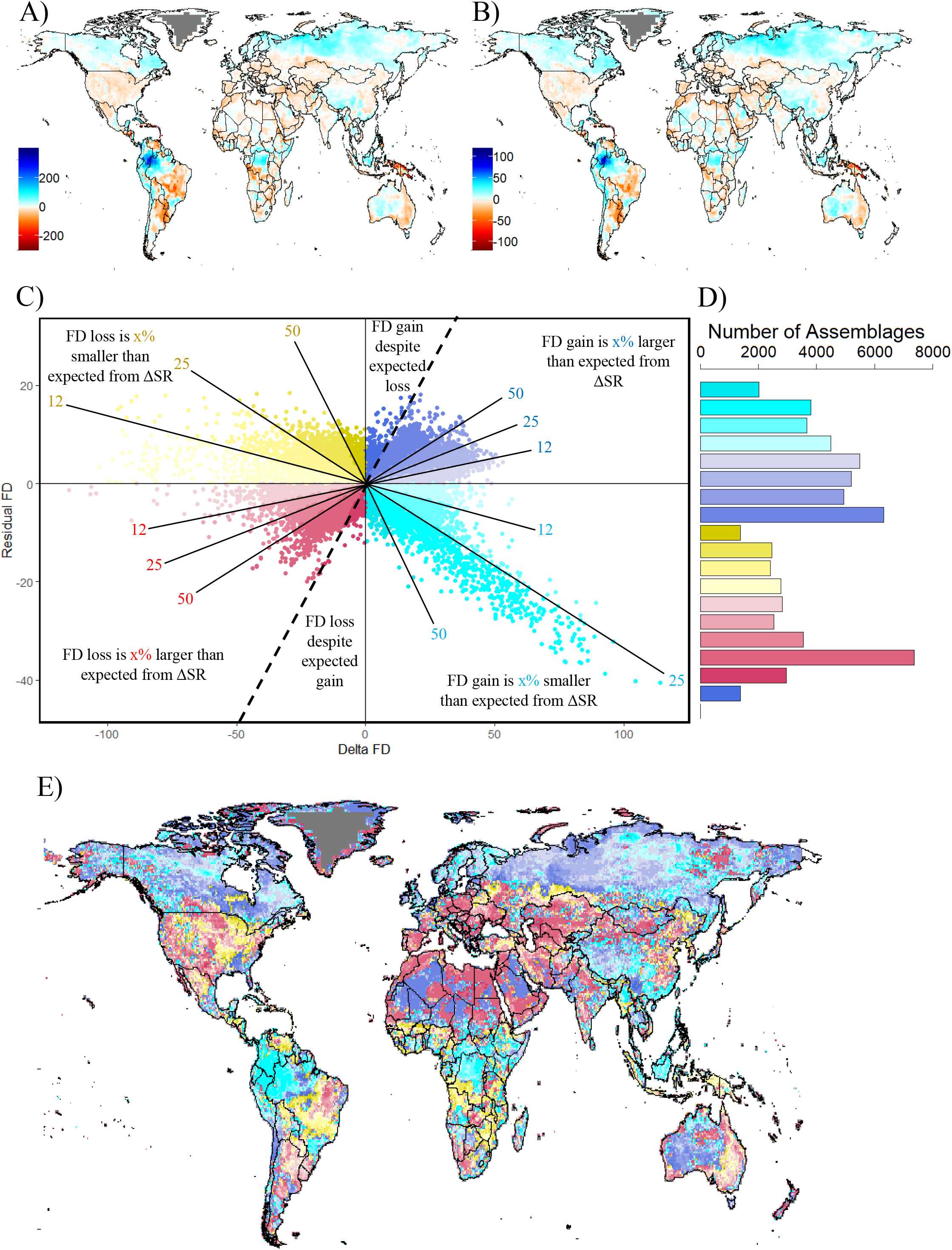
Projected cladewide changes in avian species richness (SR) and functional diversity (FD) from 1995 to 2050 under the RCP6.0 emissions scenario, under an intermediate dispersal scenario. **A)** Absolute change in SR; **B)** Absolute change in FD; **C)** The relationship between observed ΔFD and Residual FD (the difference between observed ΔFD and ΔFD predicted from ΔSR). Annotations denote regions where ΔFD is larger or smaller than predicted from ΔSR, indicating assemblages where ΔFD is driven by the gain or loss of functionally distinct and functionally redundant species respectively. Dashed line separates assemblages where ΔSR predicts FD loss (left) and FD gain (right); **D)** The number of assemblages belonging to each scenario presented in C; **E)** Geographical distribution of the scenarios presented in C. Dark grey areas in all maps indicate regions for which no data were available

Our projections showed substantial changes in all functional diversity metrics, with the direction and magnitude of effects varying with geographical location (Figure 1b). Higher latitudes showed widespread, consistent increases in FD and Hvol (Figure 1b, S2a). FD and Hvol also increased substantially in north-west South America, and to a lesser degree in inland western Australia, Tasmania, and across parts of Africa and south-east Asia. Major FD and Hvol losses occurred in New Guinea, the Caribbean, Eastern South America, and Eastern Australia. Smaller, yet consistent FD and Hvol decreases occurred in the USA, Northern Mexico, continental Europe, and the southern Congo forest-savannah.

Gains in FD were generally greater than predicted from ΔSR at higher latitudes, as well as western Australia and the Tibetan plateau (Figure 1e, dark blue regions), indicating that ΔFD in these regions was primarily driven by the gain of functionally distinct species. FD losses were greater than predicted in much of Europe and western Asia, north Africa and the Middle East (Figure 1e, red regions), indicating losses of functionally distinct species. These losses were mostly over 50% greater than predicted from ΔSR (Figure 1c,d). In contrast, FD losses which were lower than predicted, and thus driven by losses in functionally redundant species, were present in relatively few assemblages (Figure 1d), largely in Brazil, the southern Congo forest-savannah, the eastern USA, and parts of Indonesia (Figure 1e, yellow regions). However, observed ΔFD in these regions was generally larger than in regions where ΔFD was primarily driven by the loss of functionally distinct species (Figure 1c). Assemblages in which FD gains were smaller than predicted, and thus driven by the gain of functionally redundant species, were slightly less frequent than assemblages where FD gains were greater than predicted (Figure 1d) but were associated with larger observed FD increases, particularly in north-west South America (Figure 1c).

We found that FRic increased in few assemblages, mainly in parts of South America, western Australia, and northern areas of Russia and eastern Canada (Figure S2b). FRic losses were more widespread in eastern South America, the Caribbean, Venezuela, and New Guinea. Extreme FRic decreases occurred in parts of New Zealand, but were confined to only a handful of localised assemblages. Finally, the greatest Euclidean distances between baseline and projected Hcent occurred in the Sahara, the Arabian Peninsula, Northern Canada, and Greenland (Figure S2c). Overall, global mapping of all birds showed a complex mosaic of climate change effects on functional diversity, with uncertain implications for ecological processes.

### b. Frugivores

To explore the implications of climate change for seed dispersal we assessed how species richness changed for a baseline dataset of 898 frugivorous species in 37924 grid-cell assemblages worldwide. We found that frugivore assemblages are most species-rich in the tropics, with a steep decline in diversity at higher latitudes, in both the current (baseline) and future (projected) scenarios (Figure S1b,c). According to projected frugivore distributions, 1972 (5.2%) of these assemblages lose all frugivore species, and 17 species have no suitable projected climate within their dispersal buffer and are thus considered to be extinct by 2050. Conversely, 3064 previously unoccupied assemblages become occupied with at least one frugivore species. Our projected future assemblages contain 881 frugivore species in 39016 grid-cell assemblages – a net gain of 1092 grid-cells (+2.9% relative to baseline). Changes in species richness ranged from losses of 54 species to gains of 72 species (mean = 0.03, sd = 5.83).

Shifts in the trait structure of frugivore assemblages were not associated with latitude, as both gains and losses of FD occurred patchily across the tropics. The greatest frugivore FD and Hvol losses are projected to occur in New Guinea, Eastern South America, the Caribbean and Central America, and to a lesser degree in the southern Congo forest-savannah (Figure 2a,b). In contrast, gains in FD and Hvol are projected to occur in the north-west of South America, as well as regions surrounding the Gulf of Mexico and patches of South-East Asia.

**Figure 2.**
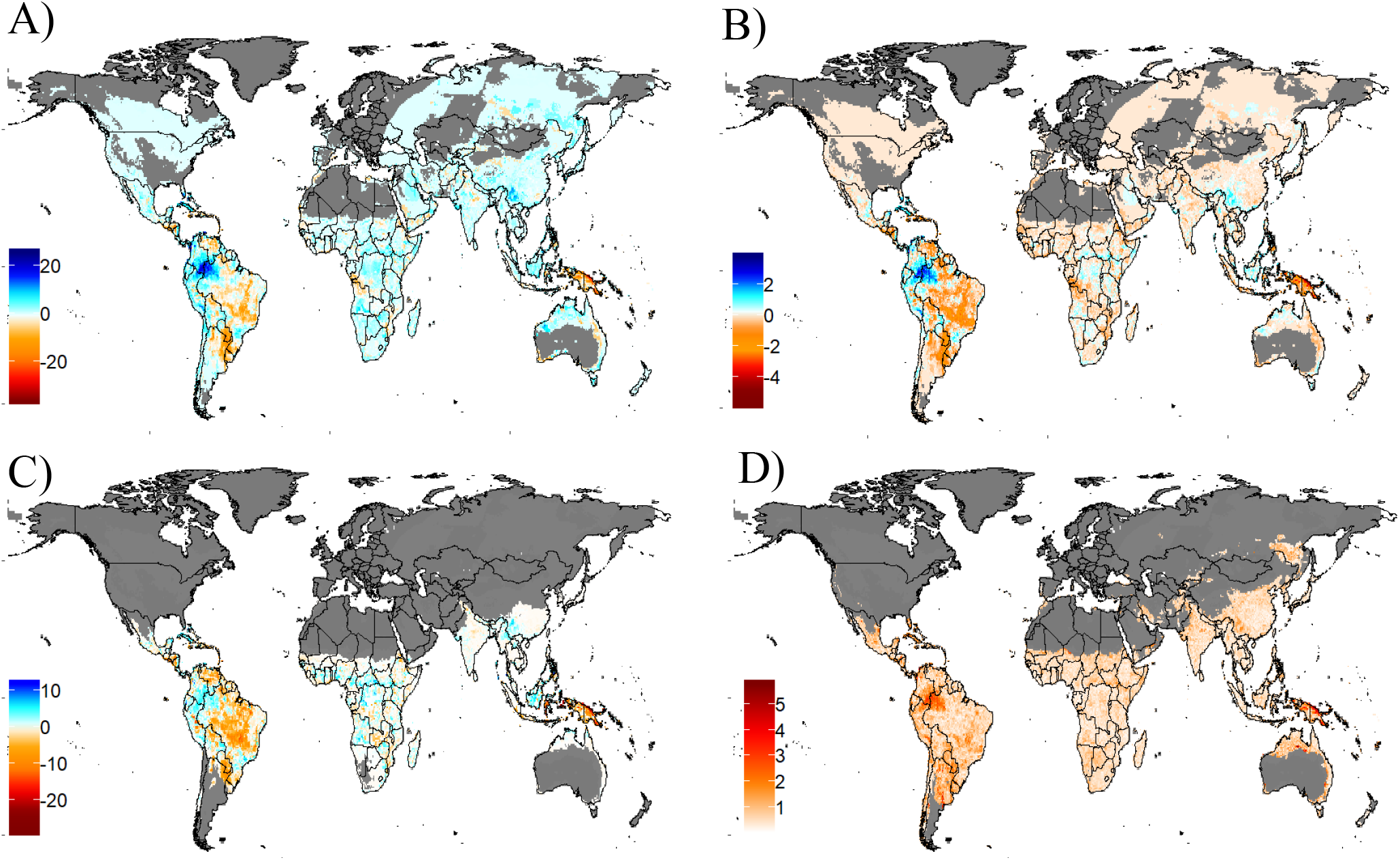
Projected **c**hanges in avian specialist frugivore functional diversity from 1995 to 2050 under the RCP6.0 emissions scenario, under an intermediate dispersal scenario. **A)** Functional diversity (FD); **B)** Hypervolume volume (Hvol); **C)** Functional richness (FRic); and **D)** Euclidean distance between current and predicted hypervolume centroids. Dark grey areas indicate assemblages for which no data were available.

In most assemblages, ΔFD was more extreme than predicted by ΔSR (Figure 3b), indicating that changes in frugivore FD tended to be driven by the gain or loss of functionally distinct species. This result was more pronounced for frugivores compared with cladewide analyses. Gains in FD were greater than predicted from ΔSR in southern and central Africa, south-east Asia, the central Andes and northern Australia (Figure 3c, dark blue regions). In most cases observed FD gains were over 50%, and in some cases over 100%, larger than predicted (Figure 3a,b). Losses in FD were greater than predicted from ΔSR in eastern South America, eastern Australia, New Guinea, and patches of Sub-Saharan Africa and Arabia (Figure 3c, red regions). The majority of these losses were in excess of 50% greater than predicted from ΔSR (Figure 3b). Furthermore, the observed FD losses in these regions were greater than those observed in any other regions (Figure 3a). We detected relatively few assemblages with lower FD losses than predicted by ΔSR. These were only associated with comparatively small changes in observed ΔFD (Figure 3c, yellow regions). Regions which gained less FD than predicted by ΔSR were also comparatively rare (Figure 3a,b), most notably occurring in the north-west of South America (Figure 3c, turquoise regions). Additionally, in many assemblages, especially beyond the tropics, no change in assemblage composition occurred (Figure 3c, light grey regions).

**Figure 3.**
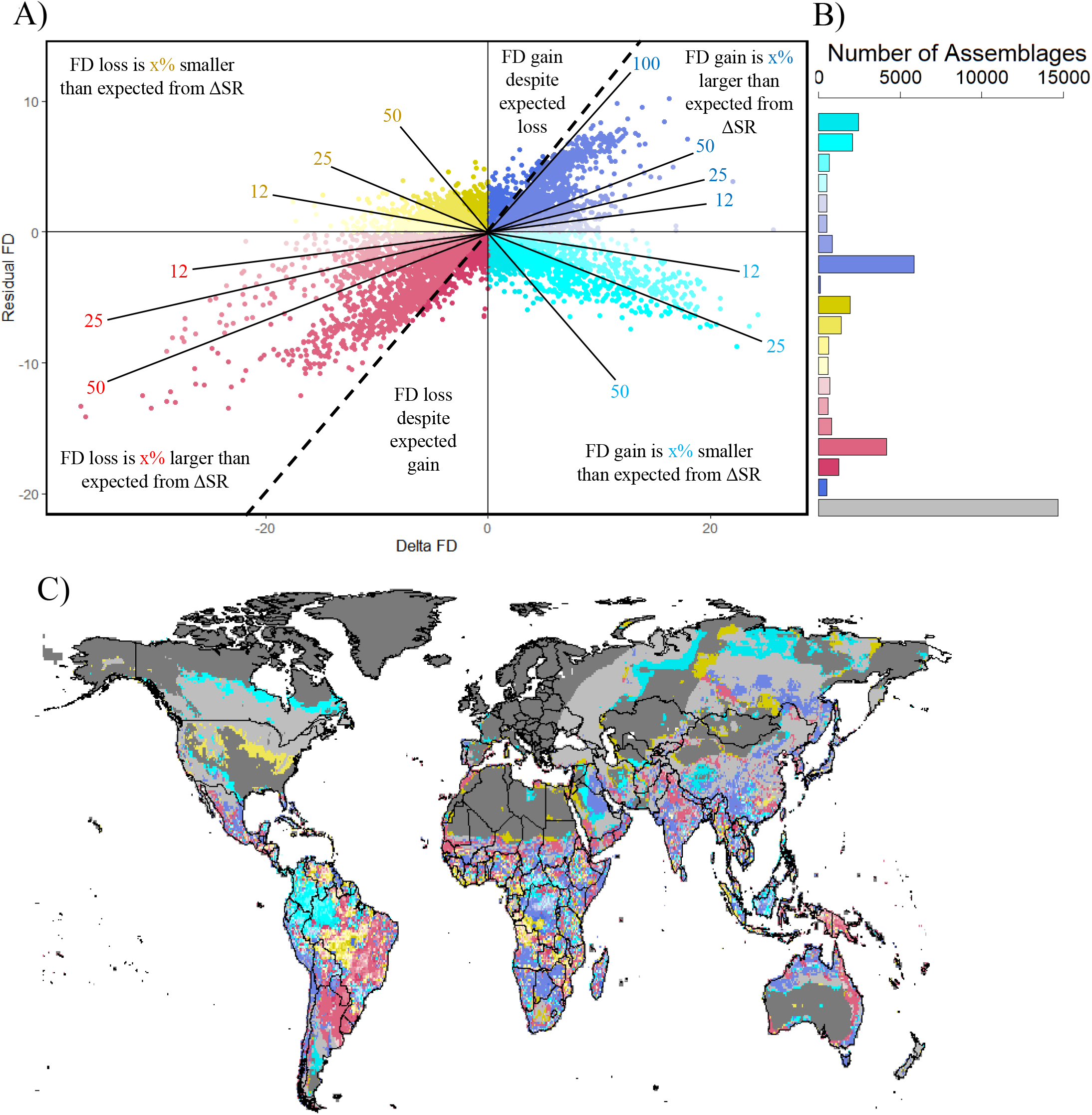
The relationship between ΔSR and ΔFD for specialist frugivores. **A)** The relationship between observed ΔFD and Residual FD (the difference between observed ΔFD and ΔFD predicted from ΔSR). Annotations denote regions where ΔFD is larger or smaller than predicted from ΔSR, indicating assemblages where ΔFD is driven by the gain or loss of functionally distinct and functionally redundant species respectively. Dashed line separates assemblages where ΔSR predicts FD loss (left) and FD gain (right). **B)** The number of assemblages belonging to each scenario presented in A. **C)** Geographical distribution of the scenarios presented in A. Light grey areas indicate regions in which no change in assemblage composition was observed. Dark grey areas indicate regions for which no data were available.

Widespread decreases in FRic occurred in South America and the Caribbean, as well as in Papua New Guinea and the eastern part of Indonesia (Figure 2c). FRic increases occurredd in north-west South America, west Africa, and Borneo. Analyses of FRic were constrained by low richness of frugivores outside the tropics: 22494 baseline assemblages and 22547 future assemblages contained fewer than five species, resulting in 23005 assemblages – most located in North America and Asia – for which ΔFRic could not be calculated. Finally, the Euclidean distance between current and projected Hcent was greatest in north-westernSouth America and in New Guinea, with other localised shifts in patches around the Gulf of Mexico and in north-eastern Australia (Figure 2d).

### c. Invertivores

Terrestrial invertivores are extremely widespread in distribution, occurring in all 65546 baseline and future grid-cell assemblages. Geographical range projections revealed that 41 (0.9%) of 4347 invertivore species in baseline assemblages had no grid-cells with suitable projected climate within their dispersal buffer and were thus assumed to become extinct by 2050. Many surviving invertivores shifted in distribution, driving changes in species richness ranging from losses of 145 species to gains of 194 species (mean = 1.63, sd = 16.85; Figure S1d, S1e).

In comparison with frugivores, the response of invertivore assemblages to climate change was far less patchy. Invertivore FD and Hvol underwent substantial and widespread increases across higher latitudes in the northern hemisphere, with losses at lower latitudes in the USA, Europe and Asia (Figure 4a, 4b). Outcomes in tropical regions were slightly less regular, with further large increases of FD and Hvol in north-west South America offset by losses in eastern South America, New Guinea, and the southern Congo forest-savannah.

**Figure 4.**
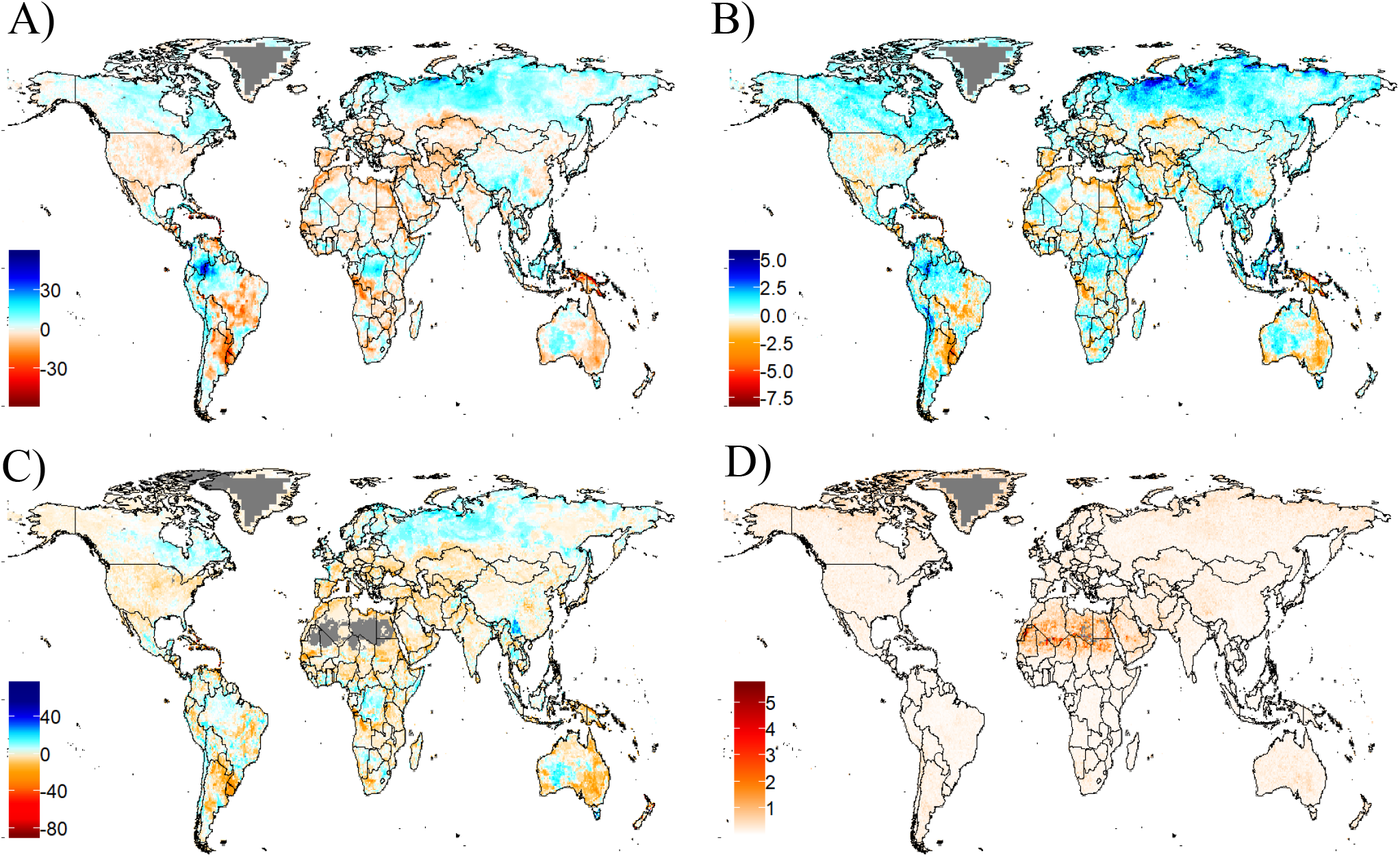
Projected **c**hanges in avian specialist invertivore functional diversity from 1995 to 2050 under the RCP6.0 emissions scenario, under an intermediate dispersal scenario. **A)** Functional diversity (FD); **B)** Hypervolume volume (Hvol); **C)** Functional richness (FRic); and **D)** Euclidean distance between current and predicted hypervolume centroids. Dark grey areas indicate assemblages for which no data were available.

Changes in invertivore FD were greater than predicted from ΔFD, especially for FD losses, and were thus primarily driven by the loss or gain of functionally distinct species (dark blue and red in Figure 5b). Echoing the pattern in raw FD, relative gains in invertivore FD were generally larger than predicted at higher latitudes in the northern hemisphere (Figure 5c, dark blue regions), whereas relative losses in invertivore FD were larger than expected at lower latitudes, including much of the central USA, Europe, and western Asia, as well as north Africa, India, and Australia (Figure 5c, red regions). In most of these regions, FD losses for many assemblages were over 50% greater than predicted by ΔSR. In scattered regions elsewhere, the relative losses of FD (Figure 5c, yellow regions), or relative gains in FD (Figure 5c, turquoise regions), were smaller than expected.

**Figure 5.**
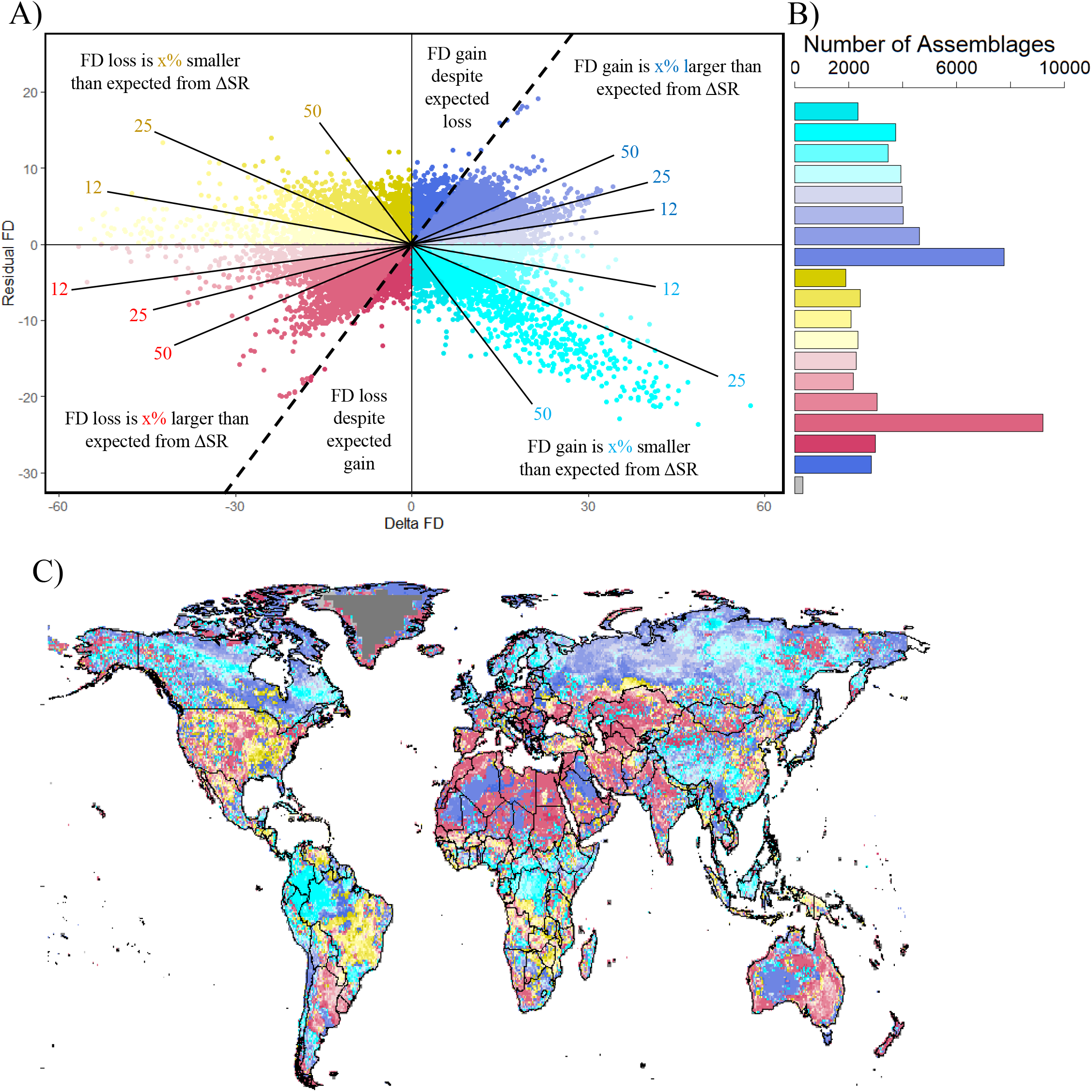
The relationship between ΔSR and ΔFD for specialist invertivores. **A)** The relationship between observed ΔFD and Residual FD (the difference between observed ΔFD and ΔFD predicted from ΔSR). Annotations denote regions where ΔFD is larger or smaller than predicted from ΔSR, indicating assemblages where ΔFD is driven by the gain or loss of functionally distinct and functionally redundant species respectively. Dashed line separates assemblages where ΔSR predicts FD loss (left) and FD gain (right). **B)** The number of assemblages belonging to each scenario presented in A. **C)** Geographical distribution of the scenarios presented in A. Light grey areas indicate regions in which no change in assemblage composition was observed. Dark grey areas indicate regions for which no data were available.

The same general pattern was observed using the FRic metric to quantify trait diversity in invertivore assemblages, with future assemblages gaining FRic mainly in patches at higher latitudes, whereas some largely tropical regions were characterised by losses in FRic (Figure 4c). The mean trait location for invertebrate assemblages was predicted to shift furthest in either arid or polar regions, with the Euclidean distance between current and projected Hcent peaking in the Sahara, the Arabian peninsula, northern Canada and Greenland (Figure 4d).

## Discussion

Our analyses show that climate change is likely to drive substantial shifts in FD for terrestrial avian assemblages across all metrics examined, with the direction and magnitude of these shifts varying by geographical location and trophic niche. In particular, our results point to a future increase in FD of invertivores at higher latitudes with concurrent losses at mid-latitudes, whereas projected shifts in FD for frugivores are more prominent in the tropics. In addition, we found that changes in FD were primarily driven by the loss or gain of functionally distinct species, particularly for frugivores. These findings suggest that the effects of climate change on future ecosystems vary across different trophic niches, and thus highlight the importance of focusing on specific ecological processes when exploring the impacts of environmental change.

When calculated for all birds, we found that shifts in trait FD occur relatively unevenly, with no clear latitudinal or geographical pattern, in line with previous analyses at the level of whole assemblages (Barbet-Massin & Jetz, 2015). Overall, we found that species richness and metrics of functional diversity tended to increase slightly at higher latitudes, presumably as a result of species ranges tracking climatic niches towards the poles (Sorte & Thompson, 2007; McQuillan & Rice, 2015). The main limitations of focusing on such generalised effects are, first, that they can mask important changes occurring within different functional groups and, second, that they cannot be interpreted in the context of specific ecological functions (Bregman et al., 2016). When we focused analyses separately on frugivores and invertivores, we found contrasting patterns of climate change impacts on these two trophic groups.

For specialist frugivores, our projections showed a mosaic of different directional effects across the tropics. Across much of South America, New Guinea, and the southern Congo forest-savannah our projections displayed a dramatic decline in frugivore FD. As specialist avian frugivores are vital dispersal agents, especially for large-fruited or large-seeded plants in South America and Australasia (Corlett, 2017; Snow, 1981), these declines in FD could threaten seed dispersal in these regions, potentially damaging biodiversity by reducing the recruitment of juvenile plants and limiting the ability of plants to track climate change (Cordeiro & Howe, 2003; McConkey et al., 2011; Mokany, Prasad, & Westcott, 2014). These declines in frugivore diversity also threaten to impair carbon storage in tropical forest ecosystems, potentially driving further climatic change (Bello et al., 2015). However, the scarcity of data on how fruiting plants will respond to climate change hinders our ability to predict exactly how ecosystem functions will be affected (Corlett, 2011).

In contrast to the tropics, many mid and high-latitude frugivore assemblages were projected to remain identical in structure under climate change. However, these regions typically contain only one or two specialist frugivore species, perhaps because dispersal by birds is predominant in the tropics (Clark, Silman, Kern, Macklin, & Islambers, 1999) with other modes of dispersal dominating at mid-to-high latitudes. Consequently, this result is probably of little importance for the provision of mid-to-high latitude seed dispersal.

Our projections for invertivores, on the other hand, show a much more distinct pattern of poleward shifts of species, particularly in the northern hemisphere. At mid-latitudes the loss of invertivores is likely to impact upon the top-down control of invertebrates, which in turn will affect other important ecosystem processes. For instance, forest productivity is likely to decline due to increased leaf damage from leaf-eating insects which are usually controlled by avian invertivores (Marquis & Whelan, 1994). Additionally, the productivity of arable farms and orchards is expected to decline due to the release of invertebrate pests from top-down control by birds (Jones, Sieving, & Jacobson, 2005; Karp et al., 2013; Mols & Visser, 2002).

Meanwhile, the ecological effects of gains in invertivore FD at higher latitudes depend largely on how invertebrate populations respond to climate change. If northward shifts are a general trend across invertebrates, the demand for top-down control resulting from shifting invertivore diversity may be matched by the supply of control from invertivorous birds, and indeed long-term studies have observed northward shifts in European odonata (Hickling, Roy, Hill, & Thomas, 2005) and lepidoptera (Parmesan et al., 1999) in response to warming over the past decades. However, other studies suggest that climate-driven changes in arthropod diversity are highly taxon and habitat-specific (Koltz, Schmidt, & Høye, 2018). For instance, Muscid flies, one of the most abundant and diverse groups of Arctic invertebrates, declined in abundance by 80% between 1996 and 2014 within Greenland (Buddle & Schmidt, 2018). Furthermore, invertebrate declines may be exacerbated by a reduction in flowering season arising from phenological differences between plant species in different areas (Høye, Post, Schmidt, Trøjelsgaard, & Forchhammer, 2013). Together, these results suggest that rather than simple changes in abundance, we should expect complex changes in invertebrate community structure to occur (Callaghan et al., 2004), suggesting that some regions will experience trophic mismatches in which the supply of top-down control from invertivores differs greatly from invertebrate abundance, potentially impairing ecosystem functions (Durant, Hjermann, Ottersen, & Stenseth, 2007; Schweiger, Settele, Schweiger, O., Settele, J., Kudrna, Klotz, & Kühn, 2008).

The results of our cladewide projections are very similar to, and almost certainly driven by, the projections for invertivores. This is unsurprising, as invertivores constitute over half of our species sample. The trends in frugivore FD, and potentially trends for other functional groups, merely dampen the observed effects at a general level. Consequently, the implications of climate change for ecological functions are best determined by studying the effects at the level of each functional group.

Some trends were, however, consistently observed across all groups. For instance, both frugivore and invertivore projections displayed a major increase in functional diversity in north-western South America, centred approximately on the Colombia-Brazil border. This increase was less than expected from the increase in species richness, indicating that the trend is primarily driven by the addition of functionally redundant species with relatively similar traits. This raises the question of whether so many functionally similar species will be able to coexist within this novel assemblage. Coexistence between species is facilitated by divergence in functional traits and increasing phylogenetic distance (Barnagaud et al., 2014; Pigot, Jetz, Sheard, & Tobias, 2018; Pigot & Tobias, 2013), so coexistence between functionally similar, closely-related species may be impossible – a “forbidden combination” which must result in the local extinction of one species (Diamond, 1975). Even where competition does not lead to local extinction it may impact species’ abundance (Fitt & Lancaster, 2017), rendering them vulnerable to extinction via stochastic causes (Lande, 1993). These impacts can further alter the functional structure of the assemblage (Mouillot et al., 2013), affecting ecosystem functions and resilience. Consequently, understanding how species interactions may structure novel assemblages under climate change is a key avenue for future research (Araújo & Luoto, 2007; Lavergne et al., 2010; Pigot & Tobias, 2013; Schleuning et al., 2016, 2020).

In our projections, changes in FD were usually greater than predicted from changes in species richness, indicating that changes in trait structure were primarily driven by the loss or gain of functionally distinct species. This contrasts with the findings of Barbet-Massin and Jetz (2015), who observed that far more assemblages, particularly in Africa and Asia, were driven by the loss of functionally redundant species. This discrepancy is likely due to differences between the trait data available: our fine-resolution continuous morphological traits enabled us to better detect functional differences between species than the relatively coarse categorical data used by Barbet-Massin and Jetz (2015).

The presence of functionally distinct species in an assemblage is vital for maintaining a breadth of ecosystem functions and services (Leitao et al., 2016), increasing the speed of ecological processes (Hedde, Bureau, Chauvat, & Deca, 2010), and promoting ecosystem stability (O’Gorman et al., 2011). As such, the loss of these species has serious effects on the ecosystem, including the unpredictable loss of function (O’Gorman et al., 2011), and so we expect that regions in which more FD was lost than predicted from species richness are likely to undergo the greatest immediate negative impacts.

In principle, the loss of functionally redundant species is less likely to have such immediate impacts, because functionally similar species still remain to maintain ecosystem functions. However, the loss of functionally redundant species still has negative effects. Functional redundancy increases ecosystem resilience (Ehrlich & Walker, 1998); loss of redundancy leaves brittle ecosystems, vulnerable to future perturbation. Our projections show that regions such as southern Africa and eastern South America, where less functional diversity was lost than predicted from species richness, may become vulnerable in this way. However, it is important to note that rapid impacts arising from the loss of functionally redundant species can still occur. There is no guarantee that one species will be able to compensate for the loss of another, for instance when one species is not abundant enough to compensate for the loss of a functionally similar species (Rosenfeld, 2002). Furthermore, other threats to avian diversity – including habitat loss, land-use change, and overhunting – may drive extinctions which leave seemingly redundant species as the sole bearers of vital ecosystem functions.

Our results differ from previous studies in that we estimate changes in FD based on more sophisticated informative continuous traits. This is an advance compared to the previously available method of calculating Gower dissimilarity between broad niche categories (e.g. Barbet-Massim & Jetz, 2015), but it relies on the availability of suitable functional trait data. It is only very recently that comprehensive functional trait datasets have become available at a global scale (e.g. Pigot et al., 2020), and these data provide excellent opportunities for furthering our understanding of the likely impacts of climate change. A particularly important avenue for future research is the consideration of species interactions (Schleuning et al., 2020). This can be accomplished by combining functional trait and phylogenetic data, as functionally similar and closely-related species are less likely to coexist (Pigot et al., 2018). Additionally, trait-based approaches may allow us to place trends in avian assemblages within the context of changes in other taxa, such as invertebrates and fruit-bearing plants, to better understand how specific ecosystem functions are likely to be affected. Currently there are considerable gaps in our understanding of how novel species combinations (e.g. frugivore-fruit combinations) are likely to function (Corlett, 2011; Høye & Culler, 2018), but functional trait data may provide insights by determining how species are likely to interact. For instance, as the dimensions of the beak are related to the size and type of food consumed (Lederer, 1975; Wheelwright, 1985), these data can be combined with data on the resource’s traits (e.g. fruit size and shape) to predict likely interactions. Such an approach has been effective in predicting frugivore-fruit interactions in Australian rainforests, especially for specialist frugivores (Moran & Catterall, 2010), and predicting the functional roles of Neotropical birds in frugivore-fruit interactions (Dehling, Jordano, Schaefer, Bohning-Gaese, & Schleuning, 2016). Advancing this line of research will allow us to predict the nature of functional changes within assemblages in far greater detail than is currently possible.

A couple of caveats must be borne in mind when interpreting our findings. Firstly, our projections are based solely on climatic data, and do not incorporate information on land-use change or other factors which may potentially drive functional diversity shifts. Our projections are also relatively coarse (grid cells represent areas over 3000 km^2^ at the equator). As species may be confined to microhabitats within each grid cell, and the climatic data represent an average over each grid cell, mismatches may occur between our projections and reality. Additionally, we used ensemble projections based on multiple models, and these models disagree to an extent about the exact responses of specific species and within specific grid cells. Furthermore, for these reasons, where our projections show no suitable range for species in the future, this should not be taken to mean that these species will certainly become extinct. Overall, while we believe our approach is robust for projecting broad macroecological trends under climate change, our findings should not be used to infer changes for specific grid-cell assemblages or individual species.

In conclusion, avian functional diversity is projected to undergo substantial, continental-scale shifts under climate change, with the direction and magnitude of trends varying with geographical location. In contrast to previous research, our analyses project that these shifts will be primarily driven by the loss or gain of functionally distinct species from assemblages. To further our understanding of how and where climate change is likely to impact ecosystem functions and resilience, future research should aim to incorporate species interactions and place changes in avian assemblages in the context of changes in other taxa, such as plants and insects. With the advent of high-quality, global-scale functional trait data, we now have a powerful tool with which to tackle these topics.

## Acknowledgements

We would like to thank Daniel Swindlehurst for assistance with the code used to calculate the functional diversity metrics. We would also like to thank Katerina Michalicckova, Bob Cregan, and Santiago Lacalle Puig for providing HPC training and technical support. CH and AV were supported by the German Federal Ministry for Education and Research (FKZ 01LS1617A), CH and MFB also acknowledges support from the Bavarian Ministry of Science and the Arts via the Bavarian Climate Research Network (bayklif) and CH also from the German Research Foundation (HO 3952/3-1).

