## Supplementary figures and tables for "Global Impacts of Climate Change on Avian Functional Diversity"

### Containing:

Table S1: Definitions of functional traits used

Table S2: Principal component scores for functional trait data

Figure S1: Projected changes in terrestrial avian species richness.

Figure S2: Projected changes in cladewide avian functional diversity metrics.

**Table S1:** Definitions of functional traits used.

| Trait | Definition |
|---|---|
| Beak length | Distance from anterior edge of nostrils to the tip of the beak |
| Beak width | Width of beak at the anterior edge of nostrils |
| Beak depth | Vertical height of beak at anterior edge of nostrils |
| Tarsus length | Distance from the notch between the tibia and tarsus to the end of the last scale of the acrotarsium |
| Secondary length | Distance from carpal joint to tip of first secondary |
| Wing Chord | Distance from carpal joint to wing tip |
| Tail Length | Distance from the tip of the longest rectrix to the point at which the two central rectrices protrude from the skin |
| Body mass | Mean species body mass |

**Table S2:** Principal Component scores for functional trait data

| | Comp.1 | Comp.2 | Comp.3 | Comp.4 | Comp.5 | Comp.6 | Comp.7 | Comp.8 |
|---|---|---|---|---|---|---|---|---|
| Standard deviation | 2.479 | 0.820 | 0.659 | 0.595 | 0.455 | 0.291 | 0.257 | 0.194 |
| Proportion of Variance | 0.768 | 0.084 | 0.054 | 0.044 | 0.026 | 0.011 | 0.008 | 0.005 |
| Cumulative Proportion | 0.768 | 0.852 | 0.906 | 0.951 | 0.976 | 0.987 | 0.995 | 1.000 |

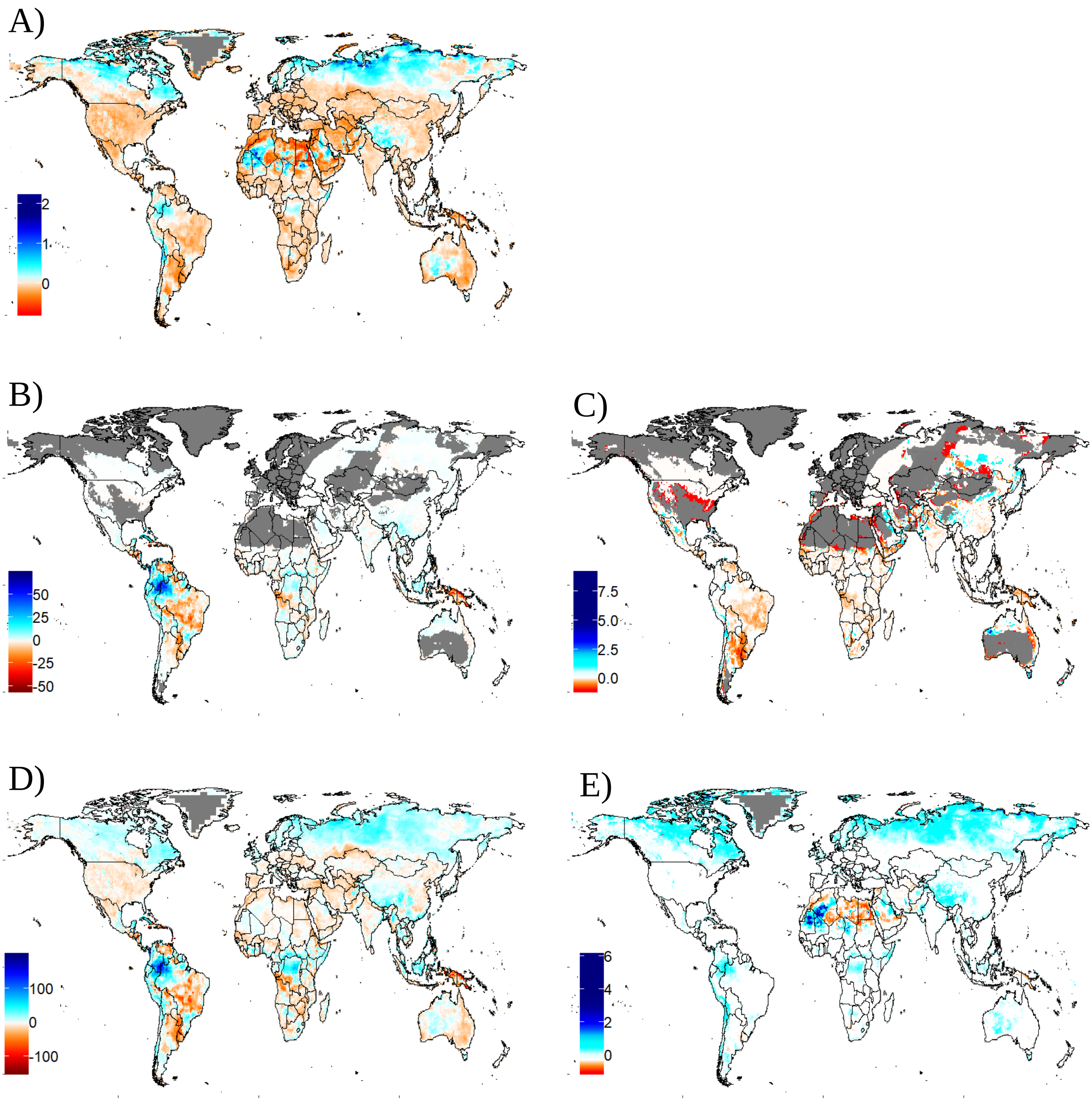


**Figure S1.** Projected **c**hanges in terrestrial avian species richness from 1995 to 2050 under the rcp6.0 climate scenario, under an intermediate dispersal scenario. **A)** Change in species richness as a proportion of baseline species richness for all bird species. **B)** Absolute change for specialist frugivores. **C)** Change in species richness as a proportion of baseline species richness for specialist frugivores. **D)** Absolute change for specialist invertivores. **E)** Change in species richness as a proportion of baseline species richness for specialist invertivores. Dark grey regions indicate areas for which no data were available. Absolute change for all bird species is given in Figure 1 (main text).

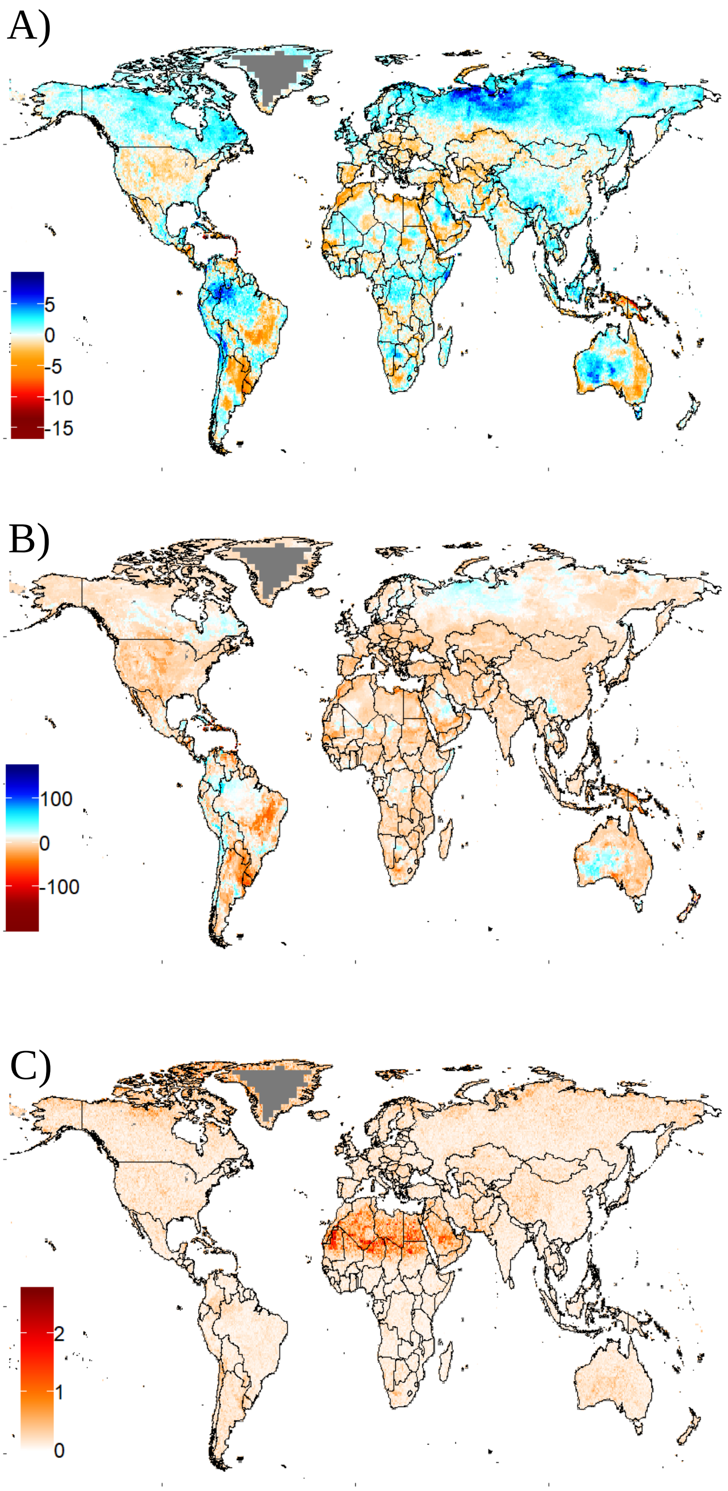


**Figure S2.** Projected changes in cladewide avian functional diversity from 1995 to 2050 under the rcp6.0 climate scenario, under an intermediate dispersal scenario. **A)** Hypervolume volume (Hvol); **B)** Functional richness (FRic); and **C)** Euclidean distance between current and projected hypervolume centroids. Dark grey areas indicate assemblages for which no data were available. Results for the Functional Diversity (FD) metric are given in Figure 1 (main text).
